# The impediment to replication at tRNA genes in *S. cerevisiae* does not require tRNA transcription, and is facilitated by topoisomerases and Rad18-dependent repair pathways

**DOI:** 10.1101/2020.01.29.925644

**Authors:** Rani Yeung, Duncan J. Smith

## Abstract

tRNA genes are widely studied sites of replication-fork pausing and genome instability in the budding yeast *Saccharomyces cerevisiae*. tRNAs are extremely highly transcribed and serve as constitutive condensin binding sites. tRNA transcription by RNA polymerase III has previously been identified as stimulating replication-fork pausing at tRNA genes, but the nature of the block to replication has not been incontrovertibly demonstrated. Here, we describe a systematic, genome-wide analysis of the contributions of candidates to replication-fork progression at tDNAs in yeast: transcription factor binding, transcription, topoisomerase activity, condensin-mediated clustering, and Rad18-dependent DNA repair. We show that an asymmetric block to replication is maintained even when tRNA transcription is abolished by depletion of one or more subunits of RNA polymerase III. By contrast, analogous depletion of the essential transcription factor TFIIIB removes the obstacle to replication. Therefore, our data suggest that the RNA polymerase III transcription complex itself represents an asymmetric obstacle to replication even in the absence of RNA synthesis. We additionally demonstrate that replication-fork progression past tRNA genes is unaffected by the global depletion of condensin from the nucleus, and can be stimulated by the removal of topoisomerases or Rad18-dependent DNA repair pathways.

## INTRODUCTION

The maintenance of eukaryotic genomes is a challenging task: replicating even the highly compact ~12MB genome of the budding yeast *Saccharomyces cerevisiae* requires the establishment and progression of hundreds of replication forks. The passage of replisomes through the genome can be impeded by a variety of protein-DNA complexes and DNA secondary structures including transcribed genes (Deshpande and Newlon 1996; Ivessa *et al.* 2003; Azvolinsky *et al.* 2009; Bermejo *et al.* 2009; Alzu *et al.* 2012; Sabouri *et al.* 2012; Osmundson *et al.* 2017; Tran *et al.* 2017), un-fired replication origins (Ivessa *et al.* 2003), centromeres (Chen *et al.* 2019), R-loops (Hamperl *et al.* 2017; Lang *et al.* 2017) and G-quadruplexes (Paeschke *et al.* 2011). Unresolved collisions between such obstacles on the DNA template and the replication fork will transiently stall or permanently arrest the replisome; this can lead to replication-fork collapse (Lambert and Carr 2013; Ait Saada *et al.* 2018), DNA damage (Ait Saada *et al.* 2018) and dissociation of the replisome from the DNA (Cortez 2015). However, widespread replication-fork stalling or arrest is detrimental even in the absence of replication-fork collapse, since the impairment of two convergent forks without a licensed origin in the intervening sequence will lead to under-replication of this region (Blow *et al.* 2011).

Transcription represents both a widespread and extensively investigated impediment to genome duplication (Hamperl and Cimprich 2016; Hamperl *et al.* 2017). Replication-transcription conflicts have been studied since the early 1980s, when it was first shown *in vitro* that DNA primer extension by the bacteriophage T4 replication machinery could be stopped by addition of RNA polymerase (Bedinger *et al.* 1983). *S. cerevisiae* tRNA genes (hereafter referred to as tDNAs) are a long-standing model system for replication-transcription conflicts. tRNAs are highly transcribed—they account for approximately 15% of cellular RNA (Warner 1999) despite representing less than 0.2% of the yeast genome. tDNAs are prominent sites of polar replication-fork pausing (Deshpande and Newlon 1996; Ivessa *et al.* 2003; Azvolinsky *et al.* 2009), whereby a tDNA oriented such that transcription occurs in the head-on orientation relative to replication impedes a greater proportion of replisomes genome-wide than tDNAs co-oriented with replication (Osmundson *et al.* 2017). Replication-fork pausing and genome instability at these sites are mitigated by the redundant activity of the Pif1-family helicases Pif1 and Rrm3 (Osmundson *et al.* 2017; Tran *et al.* 2017). Consistent with head-on replication-transcription conflicts at tDNAs being deleterious, tDNAs show a strong bias towards the co-directional orientation in the yeast genome (Osmundson *et al.* 2017). An analogous statistical co-orientation of replication and transcription is not observed for protein-coding genes in yeast (Raghuraman *et al.* 2001) but is prevalent in prokaryotes (Rocha and Danchin 2003) and has been described for both the *C. elegans* (Pourkarimi *et al.* 2016) and human replication programs (Petryk *et al.* 2016; Chen *et al.* 2019). Thus, replication-transcription conflicts can shape genome architecture and replication dynamics across biological kingdoms.

Replication-transcription conflicts at both tDNAs and protein-coding genes are associated with genome instability in yeast (Prado and Aguilera 2005; Tran *et al.* 2017). Interestingly, replisome pausing and the onset of DNA damage at tDNAs appear to be mechanistically separable: co-oriented collisions in the absence of both Pif1 helicases impede a substantial fraction of replisomes (Osmundson *et al.* 2017; Tran *et al.* 2017), but only head-on conflicts lead to a dramatic increase in R-loop-dependent gross chromosomal rearrangements (Tran *et al.* 2017). Similarly to tDNAs, transcription-dependent mitotic recombination is strongly biased to the head-on orientation at a model protein-coding gene in yeast (Prado and Aguilera 2005). In humans, codirectional and head-on conflicts with RNA polymerase II both impede the replisome, but elicit distinct signaling and damage outcomes, which are modulated by R-loop levels (Hamperl *et al.* 2017). Replication termination is enriched at the 3’-end of transcribed genes in humans (Chen *et al.* 2019), which are sites of RNAPII pausing (Glover-Cutter *et al.* 2008). This is consistent with RNAPII conflicts impeding fork movement more globally. R-loop-dependent genome instability has also been demonstrated at common fragile sites (Helmrich *et al.* 2011), and transcription-dependent instability is observed at highly transcribed genes (Barlow *et al.* 2013). Consistent with defects in RNA metabolism exacerbating genome instability in the context of a stalled replication fork, mutations that impair co-transcriptional RNA processing increase DNA damage in both yeast and humans (Paulsen *et al.* 2009; Stirling *et al.* 2012). While R-loops are associated with DNA damage at tDNAs, they do not substantially impede replisome progression at these sites (Osmundson *et al.* 2017). In prokaryotes, head-on conflicts are mutation hotspots (Srivatsan *et al.* 2010; Merrikh *et al.* 2011) and unresolved R-loops impede replisome passage at a highly transcribed head-on gene (Lang *et al.* 2017). In addition, a co-oriented conflict leads to DNA damage if RNA polymerase backtracking is increased (Dutta *et al.* 2011). Thus, there exists a complex relationship between the ultimate outcome of a conflict, the orientation of the transcribed gene, and the transcriptional state of RNA polymerase.

Although tDNAs in yeast serve as a model for eukaryotic replication-transcription conflicts, our knowledge of how these sites asymmetrically impede fork passage is incomplete. tRNAs, along with other short non-coding RNAs including 5S rRNA and U6 snRNA, are transcribed by RNAPIII. RNAPIII is a 17-subunit complex that shares some structural and functional homology with RNAPI and RNAPII (Arimbasseri *et al.* 2014). *De novo* tRNA transcription is initiated by TFIIIC binding to internal A and B box promoter elements (Orioli *et al.* 2012), which subsequently recruits TFIIIB to a position approximately 40bp upstream of the transcription start site (Paule and White 2000). Bdp1, one of three subunits of TFIIIB, facilitates the formation of a stable TFIIIB-DNA complex (Colbert and Hahn 1992; Shah *et al.* 1999; Cloutier *et al.* 2001; Kassavetis *et al.* 2005; Abascal-Palacios *et al.* 2018) resistant to dissociation by heparin and molar salt concentrations (Kassavetis *et al.* 1990). TFIIIB recruits and positions RNAPIII over the transcription initiation region, where Bdp1 stimulates allosteric rearrangement of RNAPIII subunits C37 and C34 in a manner that promotes template melting, resulting in a transcription-competent RNAPIII (Abascal-Palacios *et al.* 2018). RNAPIII recruitment to tDNAs requires all three subunits of TFIIIB (Khoo *et al.* 2014), as well as the C82-C34-C31 heterotrimeric subcomplex of RNAPIII (Brun *et al.* 1997).

Eliminating TFIIIC binding by a B-block point mutation (Baker *et al.* 1986) removed fork pausing in wild-type and *rrm3Δ* yeast (Ivessa *et al.* 2003). Furthermore, deleting terminator sequences tripled the length of the tRNA transcript (Allison 1985) without affecting replisome pausing (Ivessa *et al.* 2003). Based on these findings, it was suggested that replication fork pausing at tDNAs is caused by the transcription initiation complex (Ivessa *et al.* 2003).

tDNAs in *S. cerevisiae* are also constitutive condensin binding sites (D’Ambrosio *et al.* 2008; Haeusler *et al.* 2008). By FISH, tDNAs are sequestered in the nucleolus throughout the cell cycle, dependent on interactions between condensin subunits Smc2 and Smc4, TFIIIB subunit Brf1 and TFIIIC subunit Tfc1 (Haeusler *et al.* 2008). Depletion of the condensin subunit Brn1 leads to a decrease in intrachromosomal interactions genome-wide, and a loss of interchromosomal homotypic interactions at a subset of tDNAs (Paul *et al.* 2018). Apart from tDNAs, condensin is also enriched at centromeres and rDNA genes (Wang *et al.* 2016; Lazar-Stefanita *et al.* 2017; Schalbetter *et al.* 2017).

Unsurprisingly given their high transcriptional output (Moir and Willis 2013), tDNAs are sites of high RNAPIII transcription complex occupancy and abundant R-loops (El Hage *et al.* 2014). Convergent DNA and RNA polymerases may generate high levels of supercoiling between them, forming a topological barrier with a small footprint that makes it difficult for topoisomerases to resolve (Wang 2002) . Any combination of these factors –the transcription initiation complex, RNAPIII occupancy, R-loops and supercoiling, could contribute to the polar barrier at tDNAs. Here, we systematically deplete tDNA-associated protein complexes and assay replication-fork progression genome-wide by sequencing Okazaki fragments.

We find that transcription impedes replication-fork progression, but observe that TFIIIB creates a strong polar replication barrier even in the absence of transcription. Disruption of condensin binding does not stimulate replisome progression past tDNAs. However, impaired fork progression can be partially suppressed via inactivation of the Rad18-mediated post-replicative repair pathway or the loss of either type 1 or type 2 topoisomerase activity.

## MATERIALS AND METHODS

### Yeast strains and propagation

All strains were derived from W303 RAD5+, and most have the anchor away background: *tor1-1∷HIS3, fpr1∷NatMX4, RPL13A-2xFKBP12∷TRP1.* The exceptions are TLSΔ *rrm3Δ*, *rad18Δ rrm3Δ*, and their experimental controls (**Fig.3***rrm3Δ*, wild-type). All sequenced strains have a doxycycline-repressible DNA ligase (*cdc9∷tetO7-CDC9 cmv-LacI-URA3*). Protein tagging and gene knockout was carried out by PCR and lithium acetate transformations. Strains were grown in YPD at 30°. For spot tests, cells were grown to log phase and plated in a 1:5 dilution series. All experiments were carried out in three biological replicate strains derived from one or more crosses, except for *BRN1-FRB; rrm3Δ*, which has two biological and one technical replicate.

### Fluorescence-activated cell sorter (FACS) analysis

Cells were released and collected from G1 arrest and fixed in 70% ethanol. Fixed cells were then spun down and resuspended in 50mM sodium citrate with RNase A (Fisher 50-153-8126) for 1 hour at 50°. Samples were subsequently incubated with proteinase K (MP Biomedicals 219350480) for 1 hour at 50°, and stained with SYTOX green (Fisher S7020). Samples were sonicated before processing with a Becton Dickinson Accuri.

### Northern blots

Total RNA was extracted by phenol freeze. 10μg of total RNA was loaded onto 10% denaturing PAGE gels and run for 1h. Samples were semi-dry transferred onto zeta-probe membrane (BioRad 1620159) and blocked overnight with 125μg/mL denatured fish sperm in 6X SSPE, 0.1% SDS, 2X Denhardt’s. Membranes were subsequently probed overnight with 20pmol of radiolabeled oligos against U4 RNA (5’-CCATGAGGAGACGGTCTGG-3’) and *tS(CGA)C* (5’-TATTCCCACAGTTAACTGCGGTCAAGATATTT-3’). All blocking and probing steps were carried out at 37°C. Membranes were washed twice with 6X SSPE, 0.1% SDS before exposing to phosphoimager screens.

### Okazaki fragment enrichment and library generation

Asynchronous cultures were grown to mid-log phase, and rapamycin (1μg/mL) was added for 1h as required. DNA ligase was repressed for 2.5h with 40 μg/mL doxycycline. Cells were pelleted and Okazaki fragments were purified and deep-sequenced as previously described with some changes (Smith and Whitehouse 2012). Briefly, genomic DNA was extracted from spheroblasts using zymolyase-20T (Sunrise Science Products NC0516655) in SCE buffer (1M sorbitol, 100mM sodium citrate, 60mM EDTA, pH 7.0). Spheroblasts were resuspended in lysis buffer (50mM Tris-HCl, pH 8.0, 50mM EDTA, 100mM NaCl, 1.5% sarkosyl) and digested with proteinase K (MP Biomedicals 219350480) at 42° for 2-2.5h. DNA was precipitated and treated with rnase (RNase Cocktail Enzyme Mix, Thermo Fisher) overnight at 4° in STE. Okazaki fragments were enriched by sequential elutions from Source15Q ion exchange columns (VWR 89128-854). Up to 1μg of fragments were ligated to adaptor primer pairs overnight at 16° and cleaned (GeneJET PCR Purification, Thermo Fisher) before second-strand synthesis with Taq polymerase (NEB) at 50° for 15mins. Products between 200bp-600bp were purified from 2% agarose gels run in 0.5XTAE using QIAquick kits (Qiagen), and excess adaptors were removed by magnetic beads (HighPrep PCR, Magbio) before and after amplification (Phusion polymerase, NEB).

Sequencing was carried out on the Illumina NextSeq500 platform. All libraries are paired-end except for one biological replicate from each of: *rrm3Δ* in the anchor away background, *RPC25-FRB RPC82-FRB rrm3Δ* and *BDP1-FRB rrm3Δ*. Sequencing data for *rrm3Δ* and wild-type replicates in **Fig.4** are from Osmundson *et al.* 2017.

### Computational methods

bowtie2 (v.2.2.9) was used to align FASTQ files to the *S. cerevisiae* reference genome (SGD, R64-2-1,), and samtools (1.3.1) was used to select and sort reads with MAPQ score ≥40. BEDPE/BED files were generated using bedtools (2.26.0); and genomecov was used to calculate Watson and Crick read-depth with 1-based coordinates. The fraction of total reads mapping to either the Watson or Crick strand was calculated in 100bp bins using an in-house Python script. Rightward-moving forks were quantified as the fraction of reads mapping to the Crick strand and vice versa. Leftward and rightward-moving forks were first quantified separately, then leftward forks were flipped bioinformatically such that all forks could be represented as “rightward”-moving forks. Using R, the average fraction of “rightward”-moving forks over a 10kb window around the meta tDNA = (rev(fraction of Watson reads from leftward-moving forks) + (1−(fraction of Watson reads from rightward-moving forks)))/2. Line graphs were smoothed over 200bp. The analysis focused on fork progression around 93 tDNAs (Osmundson *et al.* 2017) that were neither near sequence gaps nor replication origins.

### Monte Carlo resampling

We evaluated the significance of the change in replication direction between datasets by Monte Carlo resampling, as previously described (Osmundson *et al.* 2017). Briefly, data across two different strains (e.g. wild-type vs. TLS*Δ*) were randomly sampled 10000 times, and the grand mean of the change in fork direction was calculated. The p-value is the number of times that the resampled data recreated a difference in means greater than or equal to the actual difference in means between the data sets.

### Data availability

Sequencing data have been submitted to the GEO under accession number GSE139860.

## RESULTS

### Quantifying replication-fork progression at tDNAs in mutant strains

The Pif1 and Rrm3 helicases were previously shown to facilitate replication-fork progression through tDNAs oriented in either direction relative to replication in yeast (Osmundson *et al.* 2017; Tran *et al.* 2017), but the precise nature of the obstacle to replication has not been clearly defined. To dissect the individual contributions of transcription *per se*, transcription factors, condensin, and DNA supercoiling to the impediment of replication-forks, we designed a panel of *S. cerevisiae* strains where potential replisome blocks at tDNAs were depleted – either by gene knock-out or using the anchor away system (Haruki *et al.* 2008) for essential genes. To elucidate the impact of TFIIIB binding and RNAPIII recruitment on fork progression, we depleted TFIIIB subunit Bdp1. We also depleted RNAPIII subunit C25 (*RPC25*), which is required for transcription initiation (Zaros and Thuriaux 2005), and subunit C82 (*RPC82*), which facilitates recruitment by TFIIIB (Brun *et al.* 1997) and tDNA promotor opening (Brun *et al.* 1997; Abascal-Palacios *et al.* 2018). Condensin subunit Brn1 was depleted to assess the effect of higher order clustering on fork progression, and topoisomerases Top1 and Top2 were depleted to determine the impact of supercoiling around replication-transcription conflicts.

All mutants were combined with deletion of *RRM3* and a repressible allele of *CDC9*, encoding DNA ligase I. The use of *rrm3Δ* as a strain background ensures that the impediment of fork progression at tDNAs is pronounced but not maximal, such that both increased and decreased fork progression should be detectable. In the presence of wild-type Rrm3, the stalling at tDNAs is sufficiently small that there are no detectable differences in fork progression upon depletion of the candidate barriers (**Fig.S1A**). The dox-repressible *CDC9* allele allows unligated Okazaki fragments to be enriched for sequencing as the basis to assay replication-fork progression.

We first validated anchor-away mediated depletion of essential proteins. Conditional depletion of essential factors recapitulated the lethal phenotypes of null mutants (**Fig. S1B**). To assay the effects of gene deletion or protein depletion on tRNA transcription, we carried out northern blots against the intron of *tS(CGA)C* (*SUP61*) such that only nascent pre-tRNAs are detected (Sethy-Coraci *et al.* 1998). Northern blotting confirmed that tRNA transcription was quickly reduced below readily detectable levels within 20 minutes of nuclear depletion of C25 or Bdp1, (**Fig. 1A**) and could be continually repressed for at least 3.5h– the maximum duration of experiments described here. Transcription shut-off in *RPC25-FRB RPC82-FRB* was equally fast (data not shown), and there was no visible difference in effect on transcription between C25 depletion alone or C25 and C82 double depletion.

**Figure 1:**
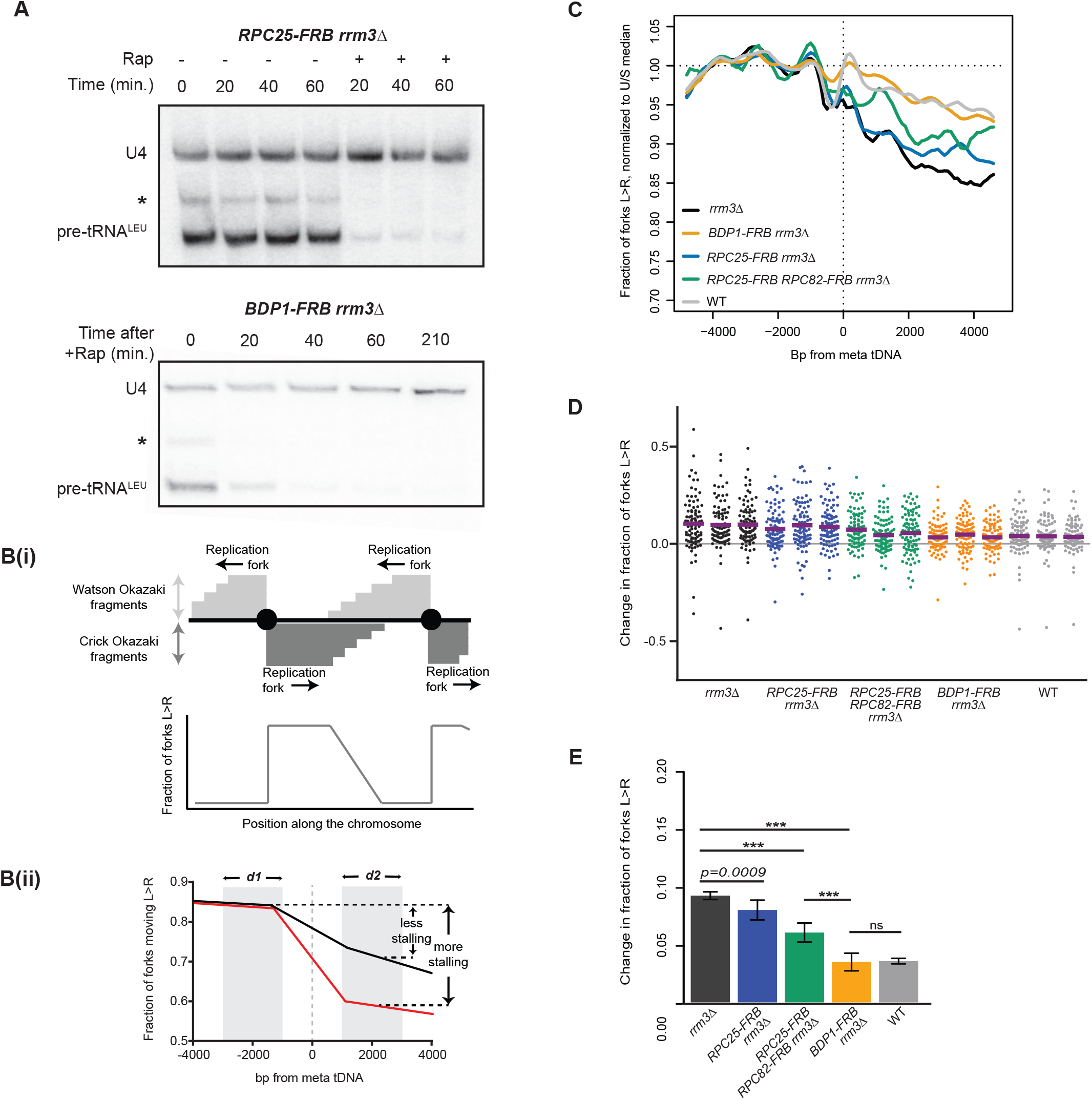
The RNAPIII transcription complex impedes the replisome at tDNAs. **(A)** Northern blots using an intronic probe against tRNA^Leu^ from *RPC25-FRB;rrm3Δ* and *BDP1-FRB*;*rrm3Δ* upon rapamycin-induced nuclear depletion of FRB-tagged proteins. * indicates a background band that we presume to be a longer pre-tRNA isoform or intermediate,10ug total RNA/lane. Loading control: U4 snRNA. **(B)** Unligated Okazaki fragments map to either the Watson or Crick strand. **(Bi)** Leftward-moving replication forks generate Watson fragments, and rightward-moving forks generate Crick fragments. (Bottom) The average direction of replication forks at every position across the genome can be quantified as the ratio of Crick:Watson reads. (**Bii**) A schematic of average fork direction ±5kb around a meta tDNA. tDNAs replicated by predominantly leftward or rightward-moving forks were analyzed separately, then superimposed such that fork direction is always presented as rightward-moving forks (Methods). Shaded areas ±1–3kb around the meta tDNA (d1,and d2) indicate the regions used to quantify the change in fork direction around the meta tDNA. **(C)** Replisome direction around tDNAs (n=93) that are at least 5kb away from origins. Values are normalized to the median fraction of rightward-moving forks upstream of the tDNA, and smoothed over 200bp. Leftward-moving forks were bioinformatically “flipped” such that they are represented as rightward-moving forks; see Methods. WT: congenic anchor away strain (Haruki *et al*., 2008). **(D)** Change in fork direction at individual tDNAs (n=93) from three biological replicates of each strain background. Mean change indicated by purple line. **(E)** Grand mean and standard deviation of the change in replication direction from above. Significance was determined by Monte Carlo resampling; *** p<0.0001.

We quantified replication-fork progression genome-wide by repressing DNA ligase (*CDC9*) for 2.5h following 1h of rapamycin treatment to deplete the protein of interest from the nucleus, and sequenced Okazaki fragments from an asynchronous population in a strand-specific manner (**Fig. 1Bi**) (Smith and Whitehouse 2012). Rapamycin treatment was maintained during ligase repression to ensure continued depletion of the protein of interest. As previously established, Okazaki fragment sequencing captures replication fork pauses that allow convergent forks to complete replication downstream of the impediment (Osmundson *et al.* 2017). Convergent forks are captured because fork direction is quantified from the strand bias of Okazaki fragments around the meta-tDNA, which includes fragments from convergent forks downstream of the tDNA. Thus, fork progression around any genomic element can be visualized as a decrease in the number of forks moving in a given direction – which is proportional to the number of Okazaki fragments mapping to a particular strand (**Fig. 1B, Methods**). This decrease can be quantified as the difference between the average number of rightward-moving forks upstream versus downstream of the meta-tDNA (**Fig. 1Bii**, d_1_ – d_2_). As previously, we used the regions ±1-3kb up- and downstream of the meta-tDNA, and focused our analysis on the 93 of 275 nuclear tDNAs in *S. cerevisiae* that are not within 5kb of an origin or sequence gap (Osmundson *et al.* 2017).

### TFIIIB (Bdp1) impedes replisome passage at tDNAs in a transcription-independent manner in *rrm3Δ* cells

In the absence of Rrm3 helicase, we observed a pronounced change in replication direction around tDNAs compared to wild-type cells, consistent with our previous analysis of these sites (Osmundson *et al.* 2017) (**Fig. 1C,**D**,**E**,**D**,**E****). Replication direction data for a meta-tDNA are shown in **Fig. 1C** for one representative replicate, and the fraction of replication forks converging within ±1 kb of each individual tDNA for each replicate is shown in **Fig. 1D**; the mean percentage of fork convergence within ±1kb of these tDNAs across replicates is shown in **Fig. 1E**. In wild-type cells with the anchor away background, 3.9% of replication termination events occur within the analyzed region, compared with 9.9% in *rrm3Δ.*

Depletion of the RNAPIII subunit C25 prior to Okazaki fragment collection in the *rrm3Δ* background moderately reduced fork progression at tDNAs, with 7.7% (p=0.0009) of forks converging at these sites (**Fig. 1E**). However, concurrent depletion of C82 and C25 yielded a significant reduction in fork stalling (6.5%, p<0.0001), though it was not a full rescue compared to the termination events in wild-type cells. Since C25 depletion abrogates detectable transcription (**Fig. 1A**), this suggests that transcribing RNAPIII acts as a barrier to fork progression but is unlikely to be the sole obstacle. By contrast, analogous nuclear depletion of the TFIIIB subunit Bdp1 dramatically reduced fork convergence at tDNAs from 10% to 3.3% (p<0.0001), essentially restoring a wild-type phenotype (**Fig. 1C,**D**,**E****). This result is consistent with the fact that TFIIIB is recruited to tDNAs before RNAPIII, and stays bound to the DNA for repeated rounds of transcription (Arimbasseri *et al.* 2014). Therefore, in the absence of C25 or a transcribing RNAPIII, fork progression is still hindered by the presence of TFIIIB and/or associated factors. The same result was obtained when all 275 nuclear tDNAs were included in the analysis (**Fig. S1C**): note that the apparent loss of strand bias upstream of the meta-tDNA is caused by the presence of proximal replication origins. For this reason, all subsequent analyses was limited to tDNAs located more than 5kb away from replication origins.

### TFIIIB (Bdp1) and RNAPIII (C25, C82) present an asymmetric obstacle to replication-fork progression in *rrm3Δ*

We and others have previously found that fork stalling at tDNAs is orientation dependent (Deshpande and Newlon 1996; Osmundson *et al.* 2017; Tran *et al.* 2017). As expected, head-on collisions were more likely to cause fork stalling than co-oriented collisions in *rrm3Δ* strains (**Fig. 2**). Bdp1 depletion in *rrm3Δ* reduced fork convergence at tDNAs from 15.1% to 5.6% (p<0.0001) for head-on collisions (**Fig. 2A,C,E**) and from 6.3% to 2.4% (p<0.0001) for co-directional collisions (**Fig. 2B,D,F**), in both cases reducing the impediment to fork progression to a level that is indistinguishable from an *RRM3* wild-type strain. Consistent with the weaker effect of RNAPIII depletion on fork progression, C82 and C25 dual depletion reduced fork stalling to 9.6% (p<0.0001) for head-on collisions and 4.3% (p<0.0001) for co-directional collisions. We conclude that the asymmetric replication-fork pausing observed at tDNAs is maintained in the absence of ongoing transcription, and that TFIIIB itself is an asymmetric impediment to fork progression in *rrm3Δ*.

**Figure 2:**
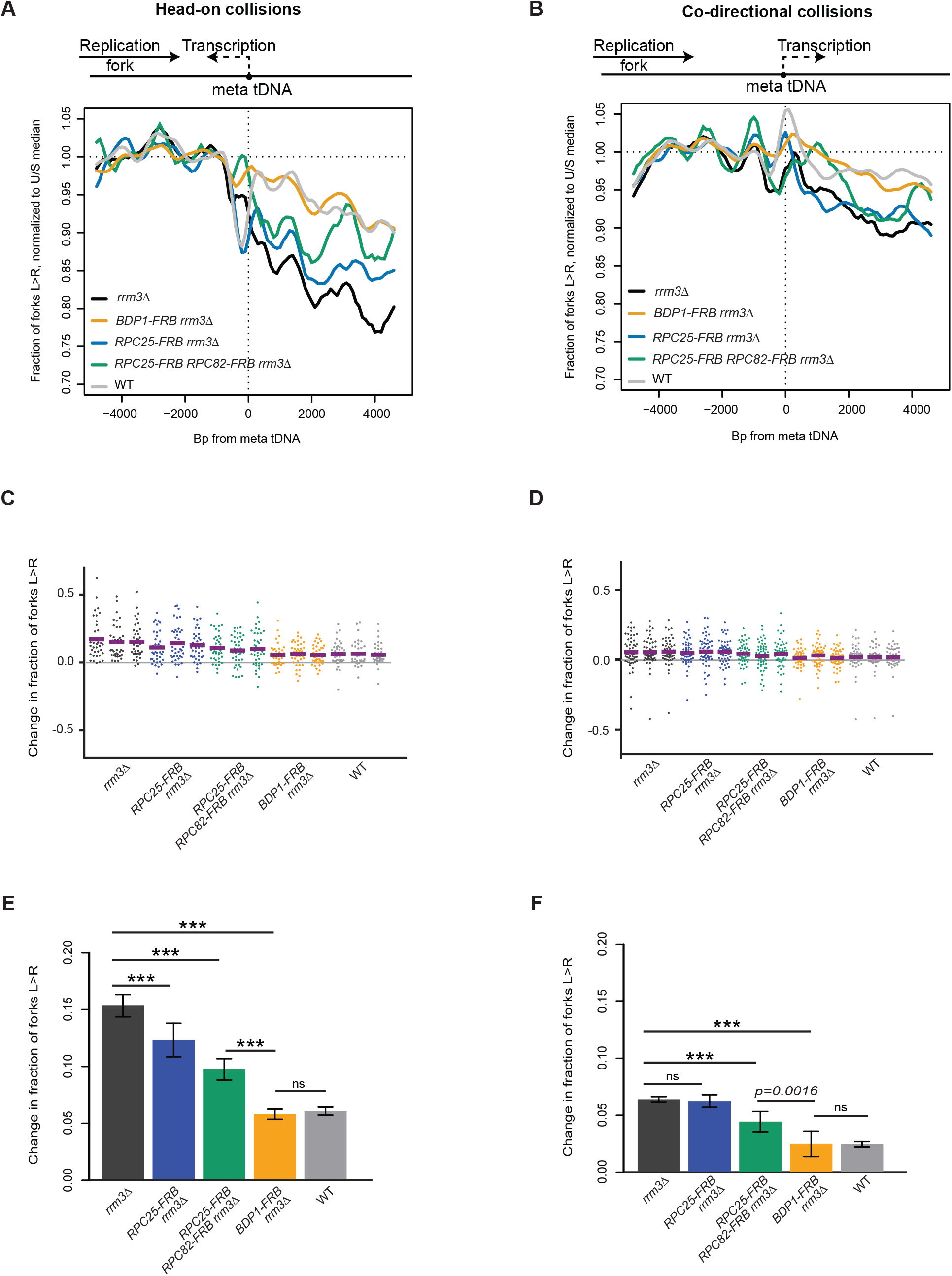
TFIIIB and RNAPIII are asymmetric obstacles to the replication fork. **(A-B)** Replisome direction around tDNAs transcribed **(A)** head-on (n=39) or **(B)** co-directionally (n=54) relative to the replication fork, and >5kb away from the closest origin. WT: congenic anchor away strain. **(C-D)** Change in fork direction around tDNAs as shown in A,B. **(E-F)** Grand mean and standard deviation of the change in replication direction from above. Significance was determined by Monte Carlo resampling; *** p<0.0001.

### Loss of condensin (Brn1) binding has no effect on replisome progression at tDNAs in *rrm3Δ* cells

tDNAs are cis-acting sites of condensin binding, and the maintenance of tDNA clustering requires direct interaction between condensin and TFIIIB/TFIIIC (D’Ambrosio *et al.* 2008; Haeusler *et al.* 2008). We depleted condensin subunit Brn1 from the nucleus to determine whether the role of condensin in DNA compaction impedes replication-fork progression at tDNAs. The Brn1-FRB construct has previously been described, and rapamycin treatment of this strain leads to dramatic loss of higher-order chromosome structure consistent with substantial loss of condensin binding (Paul *et al.* 2018). Nuclear depletion of Brn1 in the *rrm3Δ* background did not significantly rescue replication-fork progression at the 93 tDNAs analyzed, with 10.9% of forks converging at these sites compared to 10% in *rrm3Δ* alone (p=0.6713) (**Fig. 3A,D,G**). Analysis of fork progression around tDNAs separated by orientation relative to replication confirmed that condensin binding has no effect on fork pausing at tDNAs (**Fig. 3B,C,E,F,H,I)**. Fork progression did not change at the subset of tDNAs that lose interchromosomal interactions upon Brn1 depletion (**Fig. S2**) (Paul *et al.* 2018). Since a global loss of condensin binding and interchromosomal interactions does not rescue fork progression, we conclude that neither condensin itself, nor the long-range chromosomal interactions it facilitates, present a significant obstacle to replisome passage at tDNAs in *rrm3Δ*.

**Figure 3:**
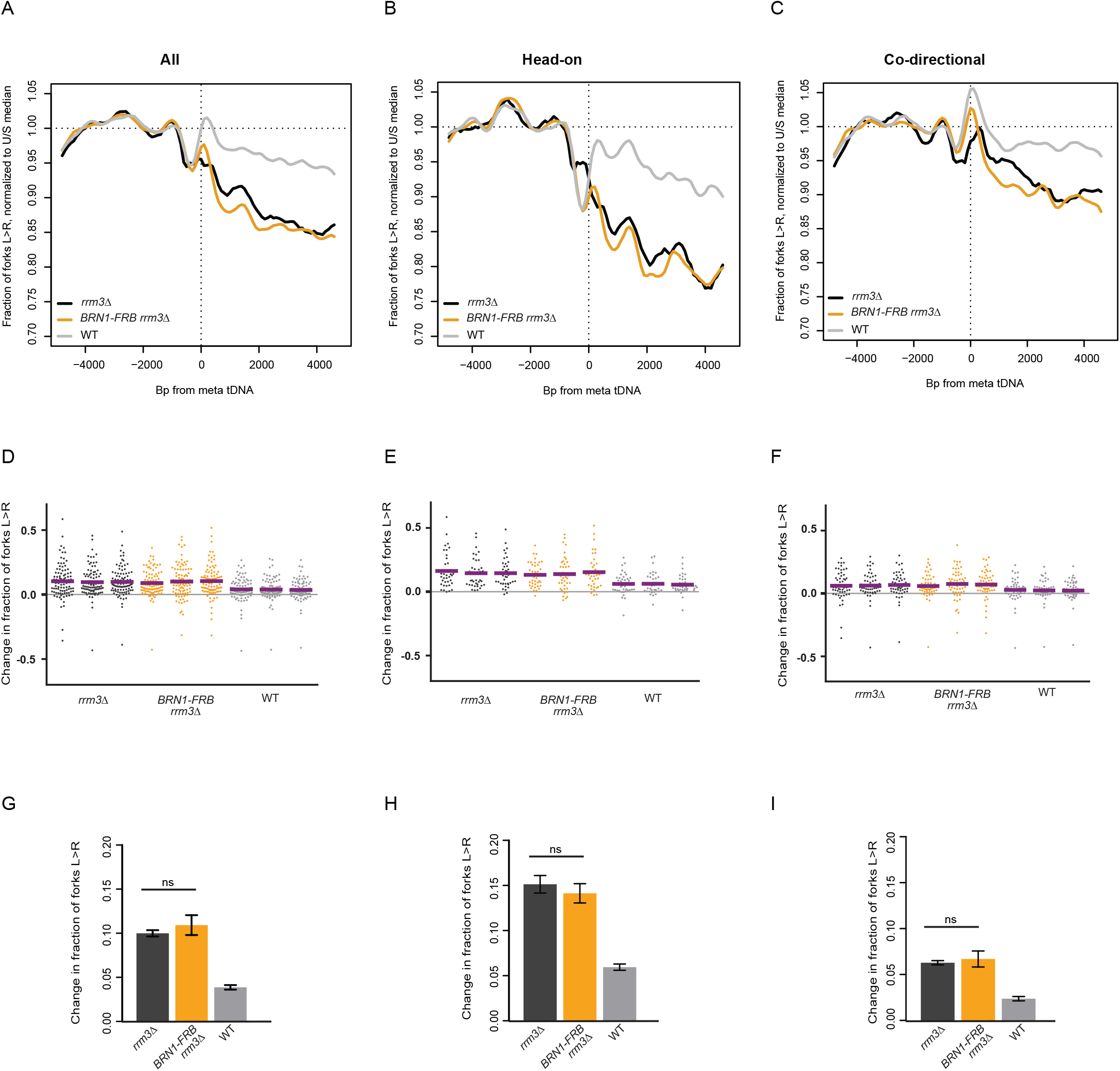
Loss of condensin binding via Brn1 depletion has no effect on replisome progression at tDNAs. **(A)** Replisome direction around tDNAs transcribed in either direction (n=93), **(B)** head-on (n=39) or **(C)** co-directional (n=54) relative to the replication fork, and >5kb away from the closest origin. WT: Dox-repressible *CDC9* strain congenic with *rrm3Δ*. (**D-F**) Change in fork direction around tDNAs transcribed in either direction (n=93), **(B)** only head-on (n=39) or **(C)** only co-directional (n=54) with respect to replication forks. Mean change from each replicate is indicated by a purple line. **(G-I)** Grand mean and standard deviation of the change in replication direction from above. Significance was determined by Monte Carlo resampling. tDNAs are >5kb away from the closest origin.

### Preventing Rad18-dependent template switching partially suppresses replication-fork pausing at tDNAs in *rrm3Δ* cells

DNA damage tolerance pathways ensure the completion of genome replication in the presence of replisome-blocking DNA damage (Cipolla *et al.* 2016). The two main pathways include error-prone lesion bypass by translesion synthesis (TLS) polymerases, and error-free template switching (TS), where the sister chromatid is used as a template to bypass fork blocks and DNA damage. Both pathways require the ubiquitylation of PCNA by E3 ligase Rad18 (Ulrich 2011). To assay the effect of DNA damage tolerance pathways on replication through tDNAs, we deleted *RAD18* and analyzed replisome progression. The absence of Rad18 in *rrm3Δ* cells significantly reduced head-on fork pausing at tDNAs from 13.2% to 10.1% (**Fig. 4B**, p<0.0001), and from 5.7% to 3.4% for co-directional collisions (**Fig. 4C**, p<0.0001). We conclude that the presence of functional DNA damage tolerance pathways impedes the replisome at *S. cerevisiae* tDNAs in *rrm3Δ*.

**Figure 4:**
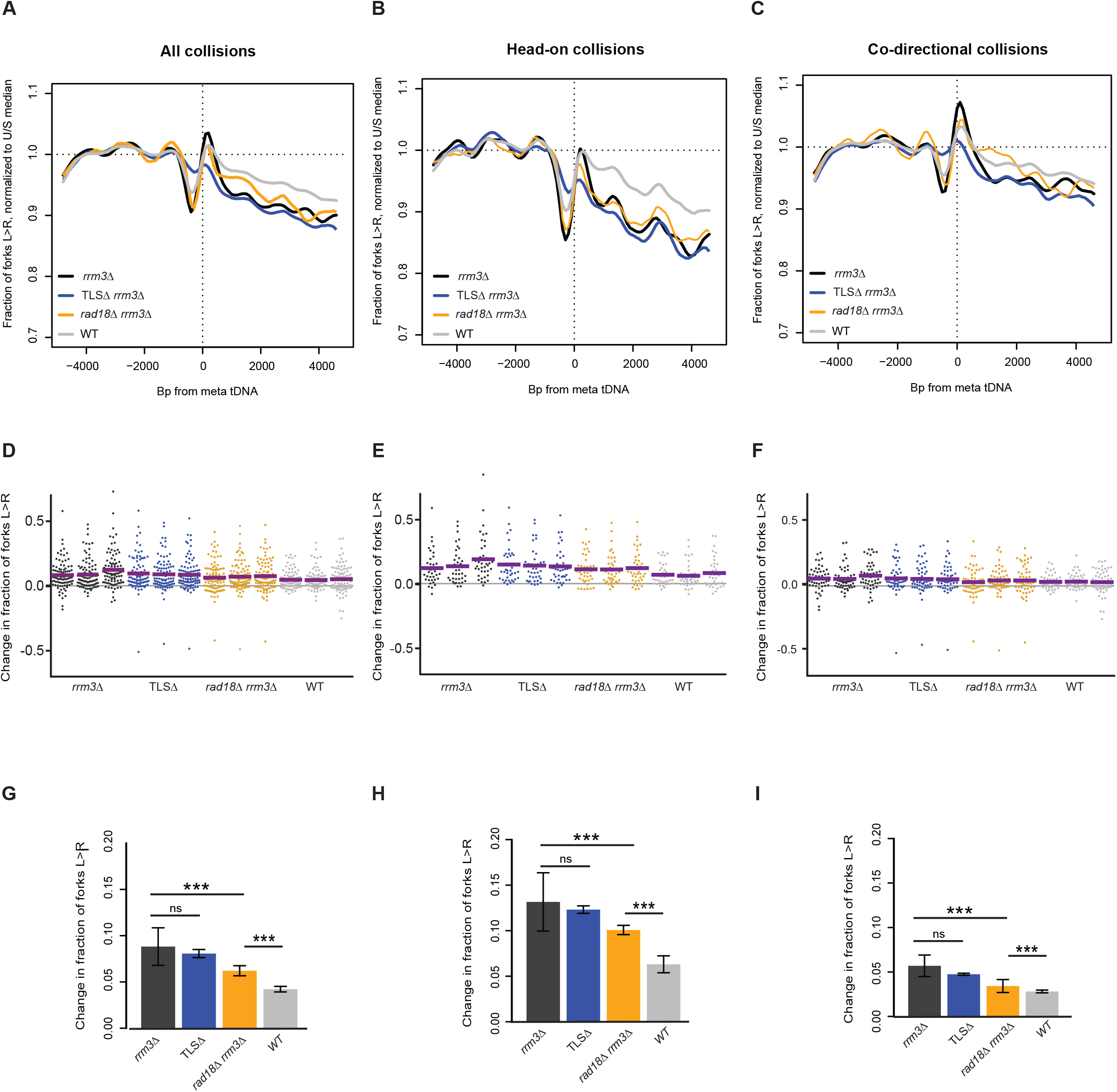
Rad18 impedes fork progression at tDNAs. **(A, D)** Replisome direction around tDNAs transcribed in either direction (n=93), **(B, E)** head-on (n=39) or **(C, F)** co-directional (n=54) relative to the replication fork, and >5kb away from the closest origin. WT: Congenic dox-repressible *CDC9* strain. Wild-type and *rrm3Δ* data are from Osmundson *et al*., 2017. (**G**) Grand mean of the change in fork direction around tDNAs transcribed in either direction (n=93), **(H)** only head-on (n=39) or **(I)** only co-directional (n=54) with respect to replication forks. Significance was determined by Monte Carlo resampling; *** p<0.0001. tDNAs are >5kb away from the closest origin. Wild-type and *rrm3Δ* sequencing data in **(A-C)** is from Osmundson *et al*., 2017.

One possible underlying cause for the increased replication-fork pausing in the presence of Rad18 could be slow replication at these sites by TLS polymerases. Even though TLS has been suggested to predominantly act on damaged DNA behind replication forks (Karras and Jentsch 2010), TLS polymerases could be recruited to stalled or paused forks (Marians 2018). To test whether any TLS polymerase was responsible for modulating replication past tDNAs, we analyzed replisome progression in a *rev1Δ rev3Δ rad30Δ rrm3Δ* strain lacking all TLS polymerases (TLSΔ). In contrast to deletion of *RAD18*, loss of TLS polymerases had no effect on replisome progression at tDNAs (**Fig. 4**). Fork progression analysis at our 93 tDNAs showed an average of 8.8% and 8.1% fork stalling in *rrm3Δ* and TLSΔ *rrm3Δ* respectively, but this difference was not statistically significant (p=0.0934). Given that TLSΔ had no effect on replisome progression at tDNAs, our data suggests that factors recruited via the Rad18 pathway for Rad5-dependent template switch may directly or indirectly hinder replisome passage through tDNAs.

### Topoisomerases 1 and 2 redundantly facilitate replication-fork pausing

The unwinding of the DNA duplex during replication creates supercoils in the unreplicated portions of the genome ahead of the fork. This mechanical strain is also transmitted behind the fork, where sister chromatids intertwine, forming precatenanes that prevent accurate chromatid segregation during mitosis. Type I topoisomerases (e.g. yeast Top1), makes a transient nick on a single strand to relax positively supercoiled DNA accumulated at unreplicated regions. Type II topoisomerases (e.g. Top2) cut both strands of DNA to resolve precatenanes behind the replication fork. DNA replication can be completed with either Top1 or Top2, but loss of Top2 leads to the formation of cruciform structures that activate the checkpoint at the M/G1 transition (Bermejo *et al.* 2007). The absence of both Top1 and Top2 can cause an accumulation of positive supercoiling (Bermejo *et al.* 2007; Yeeles *et al.* 2015; Deegan *et al.* 2019), leading to persistent Rad9-dependent checkpoint activation and ultimately S-phase arrest (Bermejo *et al.* 2007). We constructed *top1Δ rrm3Δ, TOP2-FRB rrm3Δ* and *top1Δ TOP2-FRB rrm3Δ* strains with repressible *CDC9* for Okazaki fragment analysis. Rapamycin-mediated nuclear depletion of Top2-FRB has previously been described (Bermejo *et al.* 2007), and our nuclear depletion of Top2 in a *top1Δ* strain recapitulated the expected S-phase accumulation (**Fig. S3A**). The absence of Top1 and Top2 activity did not impact tRNA transcription (**Fig. S3B**), as expected from the previously described phenotype of a *top1-1; top2-1* mutant at the restrictive temperature (Brill *et al.* 1987).

Our hypothesis was that the simultaneous loss of Top1 and Top2 would exacerbate fork stalling at tDNAs in *rrm3Δ* by increasing the buildup of supercoils adjacent to the gene. However, contrary to this expectation, the absence of topoisomerases in *rrm3Δ* significantly reduced fork stalling near tDNAs (**Fig. 5**). Only 7.8% and 7.7% of forks were stalled after Top1 deletion or Top2 depletion, respectively, compared to 10.4% in *rrm3Δ* (p<0.0001). Simultaneous Top1 and Top2 depletion further reduced fork stalling to 5.1% (p<0.0001), indicating that Top1 and Top2 act redundantly to increase rather than decrease the extent of fork stalling around tDNAs (**Fig. 5**). Interestingly, unlike the effect of Bdp1 (TFIIIB) depletion, the change in fork progression in topoisomerase-deficient cells was not precisely centered on the tDNA itself, but rather manifested more diffusely between 1kb and 5kb downstream from the tDNAs (**Fig. 5A-C,** compare to **Fig. 1C** and **Fig. 2A-B)**.

**Figure 5:**
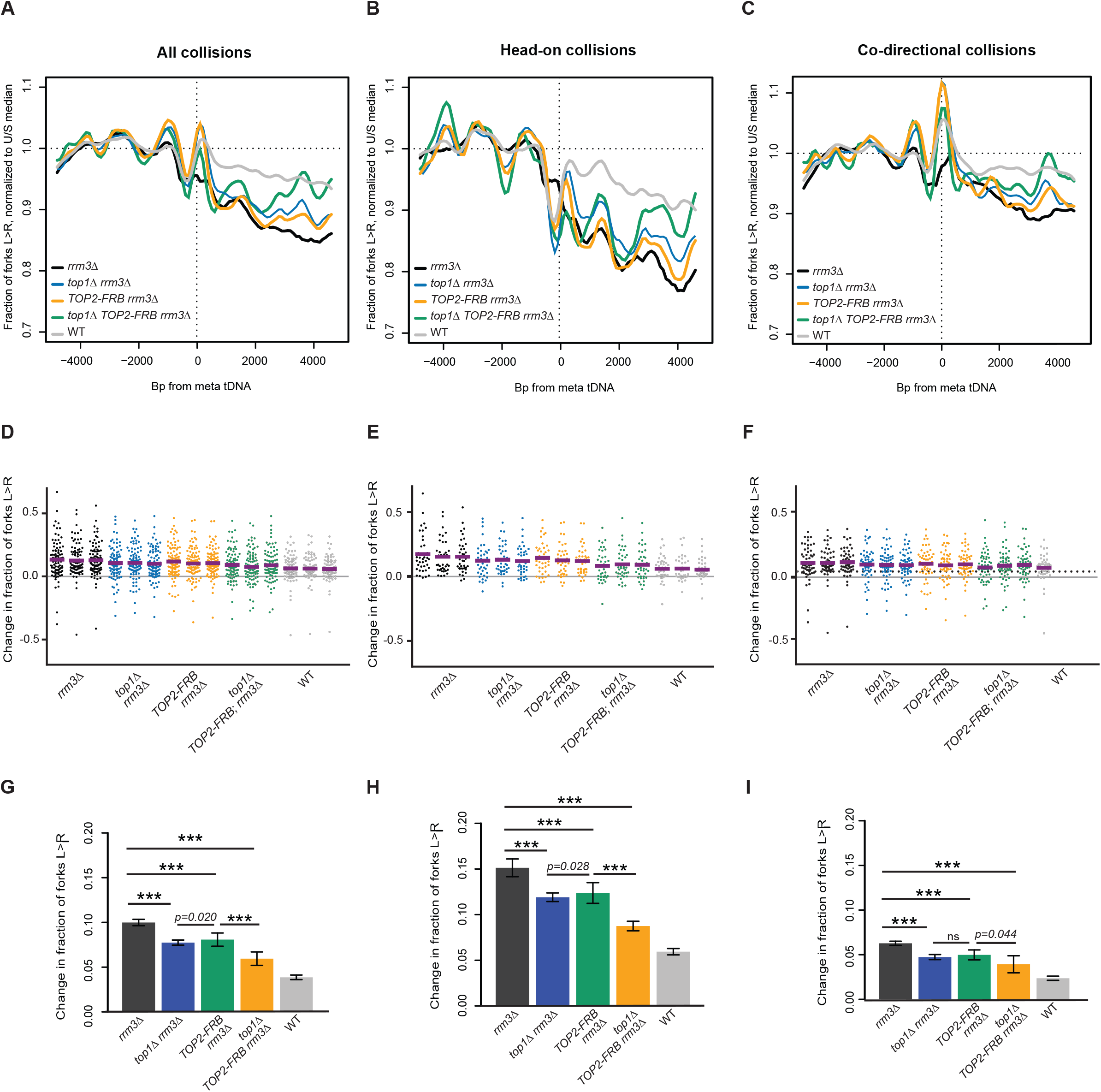
Topoisomerases facilitate fork stalling at tDNAs. **(A, D)** Replisome direction around tDNAs transcribed in either direction (n=93), **(B, E)** head-on (n=39) or **(C, F)** co-directional (n=54) relative to the replication fork, and >5kb away from the closest origin. WT: Congenic anchor away strain. Wild-type and *rrm3Δ* data are from Osmundson *et al*., 2017. (**G**) Grand mean of the change in fork direction around tDNAs transcribed in either direction (n=93), **(H)** only head-on (n=39) or **(I)** only co-directional (n=54) with respect to replication forks. Significance was determined by Monte Carlo resampling; *** p<0.0001. tDNAs are >5kb away from the closest origin. Wild-type and *rrm3Δ* sequencing data in **(A-C)** is from Osmundson *et al*., 2017.

## DISCUSSION

The progression of replication forks through a compact genome, especially at hard-to-replicate sites, is modulated by many factors. Our data indicate that TFIIIB is an asymmetric barrier to replication forks at tDNAs, whereas RNAPIII-mediated transcription is less of an impediment. We also observed that topoisomerases and Rad18 maintain fork pausing at tDNAs in wild-type cells, possibly to promote DNA repair at these sites. Replication-fork progression is the result of complex interplay between factors that cause fork stalling, and those that maintain this stalling to protect genome integrity.

### Implications for replication-transcription conflicts

When tDNAs were first identified as polar blocks to replication more than two decades ago (Deshpande and Newlon 1996), transcription by RNAPIII was determined to be integral for this impediment to replication via the use of a temperature-sensitive *rpc160* allele. It was subsequently shown by 2D gel that preventing TFIIIC binding restored fork progression at tDNAs, but transcription read-through did not affect replisome pausing (Ivessa *et al.* 2003). Our data support the idea that the most significant barrier to replisome progression is the occupancy of transcription factors, namely TFIIIB, rather than transcription.

Replication-fork progression was only slightly reduced when C25 is sequestered outside the nucleus, and still prominent when C82 is additionally depleted (**Figs. 1&2**). C25 depletion leads to impaired transcription initiation (Zaros and Thuriaux 2005), and as expected, that alone was sufficient to virtually shut off tRNA transcription within 20 minutes (**Fig. 1B**). This indicates that transcription *per se* is not required to impede the replication fork. Double depletion of C25 and C82 led to a greater reduction in fork progression, presumably because RNAPIII is not bound to TFIIIB without C82 (Brun *et al.* 1997), so the replication fork was only impeded by TFIIIB and not TFIIIB-RNAPIII. We saw the greatest rescue of fork progression when Bdp1 was depleted, leaving only TFIIIC bound at tDNAs. Since Bdp1 facilitates TFIIIB’s resistance to dissociation (Colbert and Hahn 1992; Shah *et al.* 1999; Cloutier *et al.* 2001; Kassavetis *et al.* 2005; Abascal-Palacios *et al.* 2018), we infer that TFIIIB is the primary impediment to the replication fork at tDNAs in the absence of Rrm3. TFIIIC is more labile compared to TFIIIB; tRNA transcription in *S. cerevisiae* involves extensive recycling of RNAPIII (Dieci and Sentenac 2003), mediated by the continued association of TFIIIB with the promoter after TFIIIC displacement during active transcription (Ferrari *et al.* 2004; Soragni and Kassavetis 2008; Acker *et al.* 2013). Our data suggest that TFIIIC is not a substantial impediment to the replication fork in the absence of Rrm3 (**Fig. 1C-E**).

Although tRNA transcription levels during S-phase can be modulated by Maf1 (Upadhya *et al.* 2002) and Cdk1 (Herrera *et al.* 2018), high transcription is constitutive, and maintaining tDNAs in a transcriptionally competent state is apparently sufficient to impede replication. However, because gross chromosomal rearrangements at tDNA-proximal replication forks are caused by transcription-dependent R-loops (Tran *et al.* 2017), continued TFIIIB association itself is unlikely to induce genome instability. We propose that TFIIIB occupancy has the greatest effect on replication fork progression, and RNAPIII transcription leads to genome instability.

In contrast to tDNAs in *S. cerevisiae*, which are dispersed throughout the genome but preferentially oriented in the direction of replication (Osmundson *et al.* 2017), human tDNAs are predominantly organized in large clusters with no apparent orientation bias (Mungall *et al.* 2003). *S. cerevisiae* and other unicellular eukaryotes experience strong selective pressure for increased rates of protein synthesis, while mammalian cells are subject to distinct evolutionary constraints and are therefore less likely to maximize the transcription of individual tRNAs. Therefore, these loci are unlikely to be as relevant for genome integrity in higher eukaryotes than in yeast and other unicellular models. However, replication termination in the human genome is enriched at gene 3’ ends (Chen *et al.* 2019); these sites are notable as sites of transcriptional pausing (Proudfoot 2016), which suggests that collisions between the replication fork and transcriptionally inactive polymerases are frequent in mammals – albeit with RNAPII rather than RNAPIII. It is unclear why *S. cerevisiae* RNAPII does not induce such localized replication-fork stalling: possibly the higher processivity required to allow mammalian polymerases to transcribe megabase-scale genes renders mammalian RNAPII a harder-to-displace obstacle. Regardless, because at least a fraction of relevant collisions with the replication fork appear to occur with polymerases that are not actively transcribing at the time of the collision, transcription *per se* might represent an imperfect measure of how a given locus will impact replication-fork progression.

### Replisome progression through a structured chromatin environment

tDNAs are highly enriched for condensin binding in *S. cerevisiae* (D’Ambrosio *et al.* 2008; Haeusler *et al.* 2008). The association of condensin at these sites is downstream of TFIIIC recruitment and mediates the clustering of condensin-associated tDNAs (Haeusler *et al.* 2008). tDNA clustering is lost along with other intra- and interchromosomal interactions upon condensin depletion (Paul *et al.* 2018), but this destruction of chromosome architecture does not increase the likelihood of replisome passage at tDNAs (**Fig. 3**). It is likely that the higher-order structure maintained by condensin is more labile than the protein-DNA interactions that facilitate tRNA transcription – at least during S-phase.

### Accessory factors that enhance replication-fork stalling at tDNAs

Our data demonstrate that the absence of topoisomerases significantly decreases the extent to which replication forks stall at tDNAs (**Fig. 5**). The role of topoisomerases at these loci is likely to be complex. Topoisomerase I has been shown to promote fork progression by removing R loops in mammalian cells (Tuduri *et al.* 2009) and tDNAs were sites of increased Top2 binding in a defective chromatin remodeling background in *S. cerevisiae* (Swanston *et al.* 2019). Top2 was required for the transcription of long (>3kb) transcripts by RNAPII (Joshi *et al.* 2012) in yeast, but not required for transcription of short transcripts like U4 snRNA or RNAPIII-transcribed genes (**Fig.S3B**). In support of this, inhibition of Top1 and Top2 had no effect on 5S RNA and tRNA transcription in HeLa cells *in vitro* (Gottesfeld 1986). Topoisomerases play a role in facilitating programmed fork arrest and chromatin stabilization, which allows for the binding of potential fork impediments (Teves and Henikoff 2014). The promotion of fork stalling around tDNAs by topoisomerases is supported by recent work demonstrating that topoisomerases are recruited by Tof1 to stabilize forks (Shyian *et al.* 2019), though it is unclear how the supercoiling-related activity of topoisomerases is balanced and regulated with respect to its fork stabilization activity.

Similarly to topoisomerases, loss of Rad18 decreases replication-fork pausing at tDNAs (**Fig. 4**). Deletion of all three TLS polymerases did not recapitulate this phenotype, indicating that the decreased fork progression observed in *rad18Δ* was not due to the slow activity of these polymerases. Components of the Rad5-Ubc13-Mms2-dependent template switching pathway represent plausible candidates for modulating fork stalling. This is supported by the fact that Rad5-dependent template switching suppresses duplication-mediated GCRs but not single-copy sequence mediated GCRs (Putnam et al. 2010), which are elevated in head-on conflicts at tDNAs in *rrm3Δ* (Tran *et al.* 2017). Template switching up to 75kb downstream of collapsed forks was detected in *S. pombe* (Jalan *et al.* 2019), and an alternative method of recombination-dependent fork restart – known as HoRReR – has also been reported downstream of a polar replication-fork block (Lambert *et al.* 2010; Miyabe *et al.* 2015).

Rad5 also recruits TLS polymerases to ssDNA gaps in S-phase by PCNA_K164_Ub in response to replication stress by HU treatment or in *pol32Δ* (Gallo *et al.* 2019). However, given that the TLSΔ strain had no effect on fork progression, it appears unilikely that replication-transcription conflicts at tDNAs create substrates for TLS-mediated repair. This suggests that Rad5-dependent template switching could be the primary mechanism for maintaining fork progression at replication-transcription conflicts. Slower fork progression at tDNAs may allow for the recruitment of DNA repair factors and subsequent processing of DNA damage. It will be interesting to determine the mechanism(s) by which Rad18 directly or indirectly impedes replication-fork progression at hard-to-replicate loci.

## Acknowledgements

We thank members of the Smith lab for helpful discussions and critical review of the paper. This work was funded by NIGMS R01 GM127336 to D.J.S.

---

**Figure S1:**
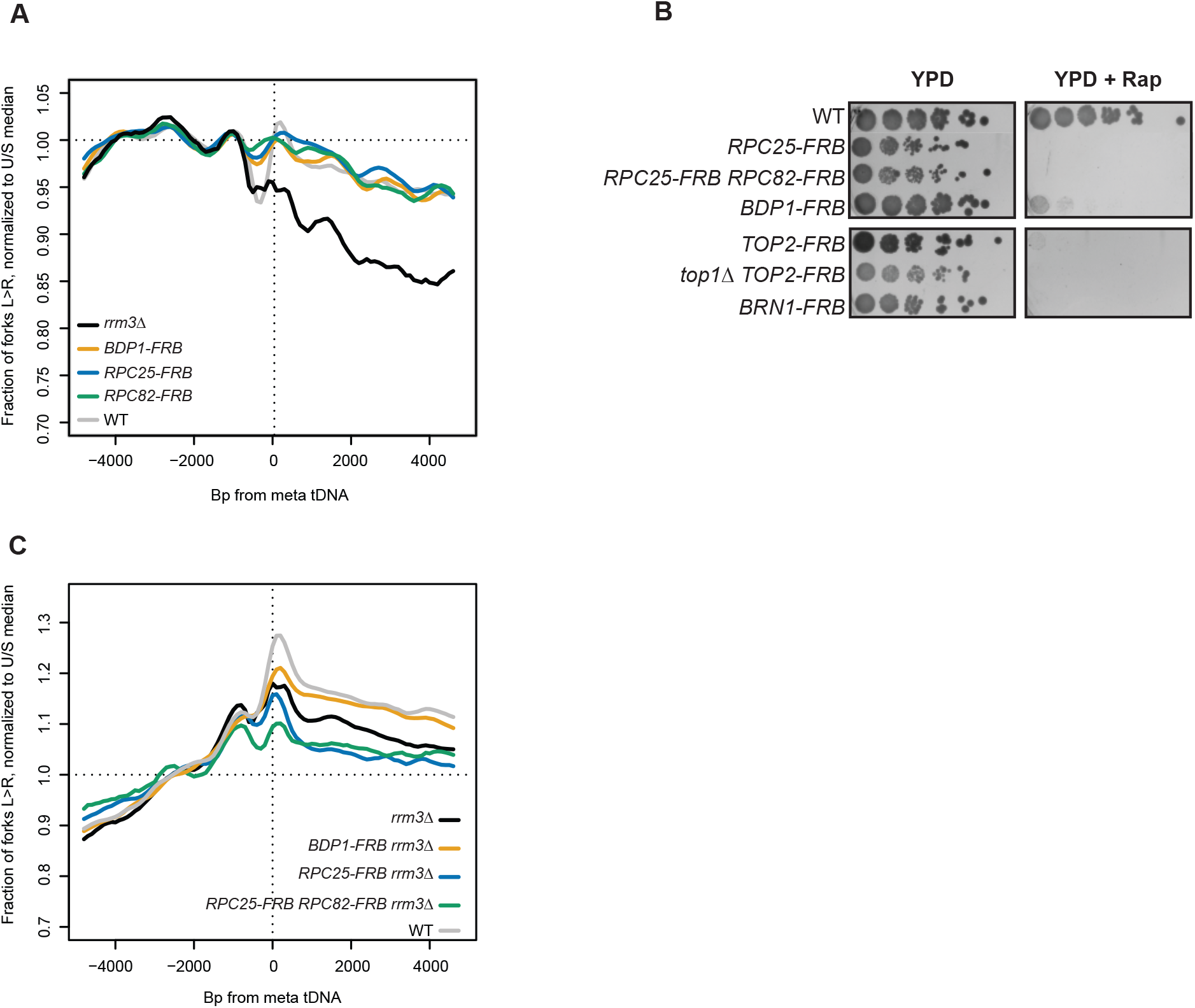
The RNAPIII transcription complex impedes the replisome at tDNAs. **(A)** Depletion of candidate factors have no effect on replisome direction with wildtype Rrm3, tDNAs >5kb from origins (n=93). The strain labeled as WT is the congenic anchor away background strain. **(B)** Serial dilutions of the indicated strains on YPD ± rapamycin at 30º. WT: congenic *rrm3Δ* anchor away strain **(C)** Replisome direction ±5kb around all 275 tDNAs. WT: congenic anchor away strain.

**Figure S2:**
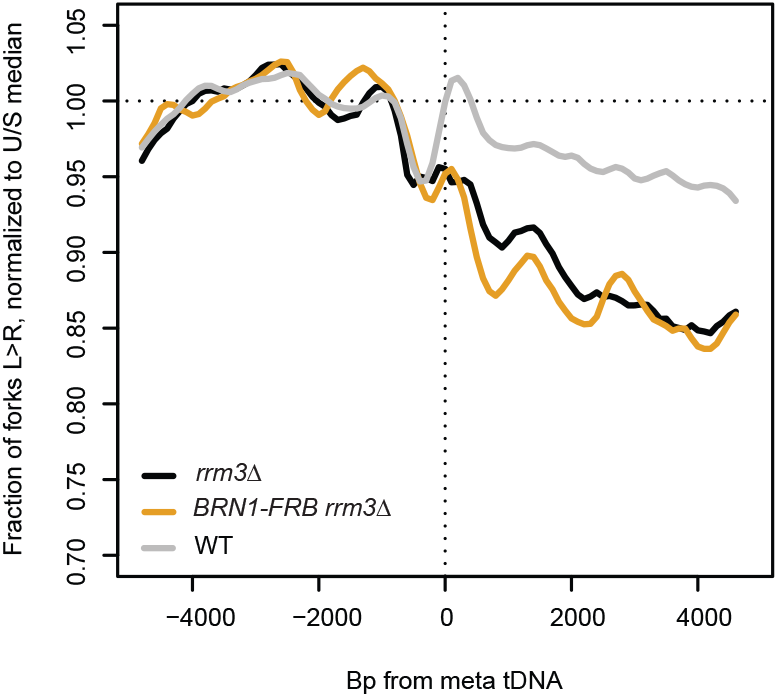
Loss of tDNA-tDNA interactions via Brn1 depletion has no effect on replisome progression at tDNAs. Replisome direction around tDNAs transcribed in either direction, >5kb away from the closest origin, that lost homotypic interchromosomal interactions after Brn1 depletion (n=25) (Paul *et al*., 2018). WT: Dox-repressible *CDC9* strain congenic with *rrm3Δ*.

**Figure S3:**
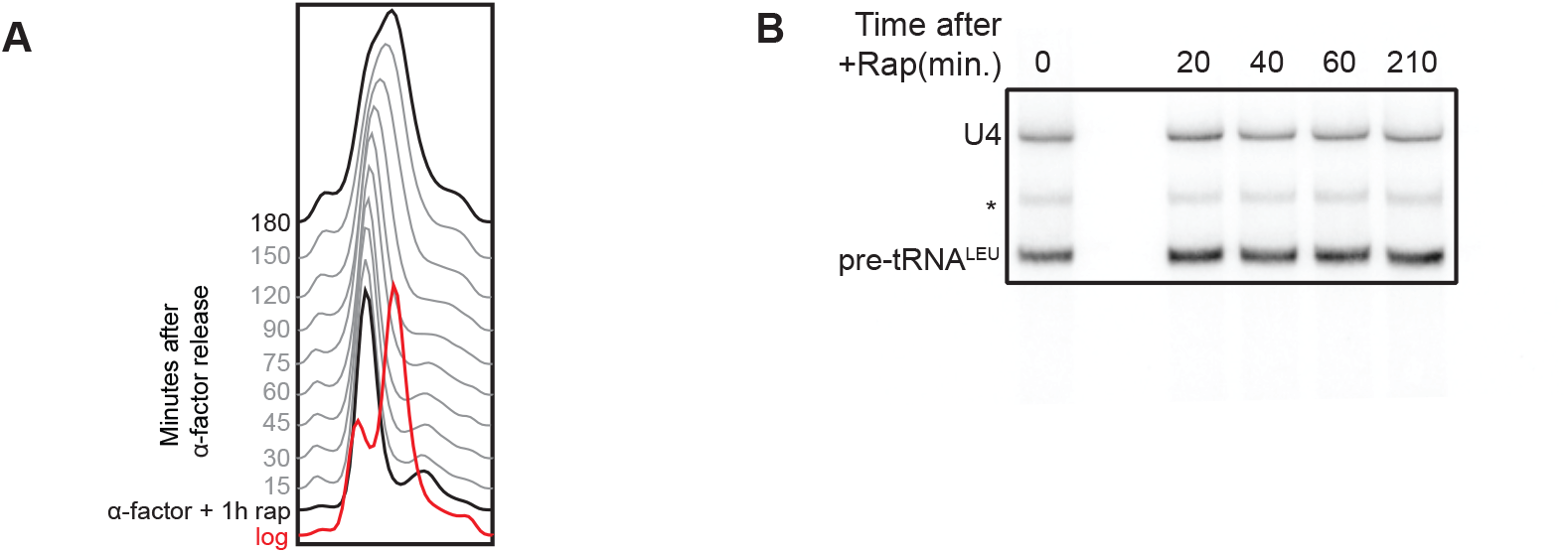
The effect of topoisomerase depletion on replication and transcription. **(A)** Flow cytometry analysis of DNA content of asynchronous *top1Δ TOP2-FRB rrm3Δ*. **(B)** Northern blot using an intronic probe against tRNA^Leu^ from *top1Δ TOP2-FRB rrm3Δ* upon rapamycin-induced nuclear depletion of FRB-tagged proteins. * indicates pre-tRNA intermediates,10ug total RNA/lane. Loading control: U4 snRNA.

